# MYC and MAX drive the reactivation of the genome after mitosis

**DOI:** 10.1101/2023.08.13.553120

**Authors:** Inma Gonzalez, Almira Chervova, Pedro Escoll, Luis Altamirano-Pacheco, Florian Mueller, Agnès Dubois, Pablo Navarro

## Abstract

Shortly after cell division, a robust wave of hyper-transcription reactivates the genome^1-3^. This phenomenon is particularly pronounced in pluripotent cells^4^, which necessitate rapid transcriptome reactivation to maintain their undifferentiated state and prevent premature differentiation. While recent work has illuminated how specific groups of genes are reactivated^4-8^, the mechanisms enabling the global, efficient and accurate post-mitotic reactivation of the genome remain unknown. Here we elucidate the direct involvement of the MYC/MAX transcription factors in the post-mitotic reactivation of pluripotent mouse embryonic stem cells. While MYC undergoes extensive phosphorylation and largely dissociates from its DNA binding sites during mitosis, we report that MAX remains bound to its targets, preferentially at promoters, and facilitates early recruitment of MYC following mitosis. Through the application of MYC/MAX heterodimerization inhibitors, we demonstrate their indispensable role in sustaining hyper-transcription in ES cells, including during the critical transition from mitosis to G1 phase. Our findings uncover a novel role for MAX in mitotic book-marking, highlighting its pivotal role in post-mitotic MYC recruitment and the re-establishment of high global transcription levels. These findings hold significant implications for medically relevant contexts, particularly when cell proliferation is of paramount importance^9^. We anticipate that the study of mitotic bookmarking by MYC and MAX and of the effects of anticancer drugs targeting MYC/MAX interactions in such process^10-12^ will be relevant for our understanding of cancer and its potential treatments.

## Introduction

The equal partition of the genetic information during mitosis is accompanied by a global shut down of transcription^**13**^. The resulting daughter cells thus need to fully reactivate their transcriptome. Several mechanisms have been proposed to enable the daughter cells to reactivate the right set of genes, including mitotic bookmarking processes whereby certain gene regulators, most notably transcription factors (TFs), remain capable of engaging in site-specific interactions with at least a subset of their targets^**14-15**^. While the number of mitotic bookmarking TFs has increased considerably over the last years, none has been shown to play a determinant role in the efficient and global reactivation of the genome following mitosis. This is particularly true in mouse pluripotent Embryonic Stem (ES) cells, where several mitotic bookmarking TFs have been recently identified but only some shown to assist gene reactivation of particular sets of genes^**4**,**7**,**16-18**^. Since these self-renewing cells display an almost inexistent G1 phase and start replication almost immediately after undergoing mitosis^**4**,**19**^, more efficient and global mechanisms are thus to be expected to drive a very rapid and accurate post-mitotic gene reactivation. Indeed, ES cells have been shown to reactivate their genome extremely rapidly after mitosis, through a burst of hyper-transcription that also takes place in other cell types but is particularly fast and global in ES cells^**1-4**^.

## Results

### MYC globally activates transcription after mitosis

We first aimed at testing whether the MYC oncogene, a potent TF involved in cell proliferation and global hyper-transcription in several contexts^**20-22**^, drives post-mitotic gene reactivation in ES cells. Indeed, its canonical binding motif, the E-box, has been previously shown to be enriched at the promoters of the most rapidly and strongly reactivated genes^**4**^. In agreement with this, we found that among the 36,385 MYC binding regions that we identified (**Table S1, Fig. 1A**), a large proportion lies close to genes (**Fig. 1B**) and overlaps with promoters (**Fig. 1C**), reproducing previous data^**23-25**^, but also with other gene regulatory elements, particularly proximal enhancers (**Fig. 1C**). Statistical analyses of the association between MYC binding sites across the genome and 5 sets of genes displaying different post-mitotic gene reactivation dynamics^**4**^ (**Table S2**), revealed a very strong link between the efficiency of gene reactivation and the presence of MYC in the vicinity of the promoter (**Fig. 1D**,**E**). The reciprocal analysis, comparing the reactivation mean of groups of genes identified by the proximity of MYC binding sites, confirmed such association (**Fig. S1A**). Thus, while the correlation between MYC binding and gene activity in interphase, which we reproduce here (**Fig. S1A**,**B**), has been established in several contexts^**20-25**^, these observations indicate a direct function of MYC in post-mitotic gene transcription. To establish this functionally, we incubated ES cells with an inhibitor of MYC binding to DNA (10058-F4, thereafter MYCi) that also leads to its partial degradation (**Fig. S1C**) and monitored nascent transcription by EU incorporation across different stages of the cell cycle (**Fig. S1D**). We found that in the presence of MYCi, the levels of global transcription were strongly reduced as early as cells undergo mitosis (early G1) although they were not completely abolished compared to those obtained upon direct transcription inhibition with flavopiridol (**Fig. 1F**). These strongly reduced levels of transcription correlate with reduced cell size (**Fig. S1E**) and affect the production of essential proteins for ES cells such as the TFs NANOG and ESRRB (**Fig. 1G, Fig. S1F**). We conclude, therefore, that MYC binds in the vicinity of the most active and most rapidly reactivated genes and directly promotes the strong and global transcriptional output characterizing ES cells as they complete mitosis.

**Fig. 1.**
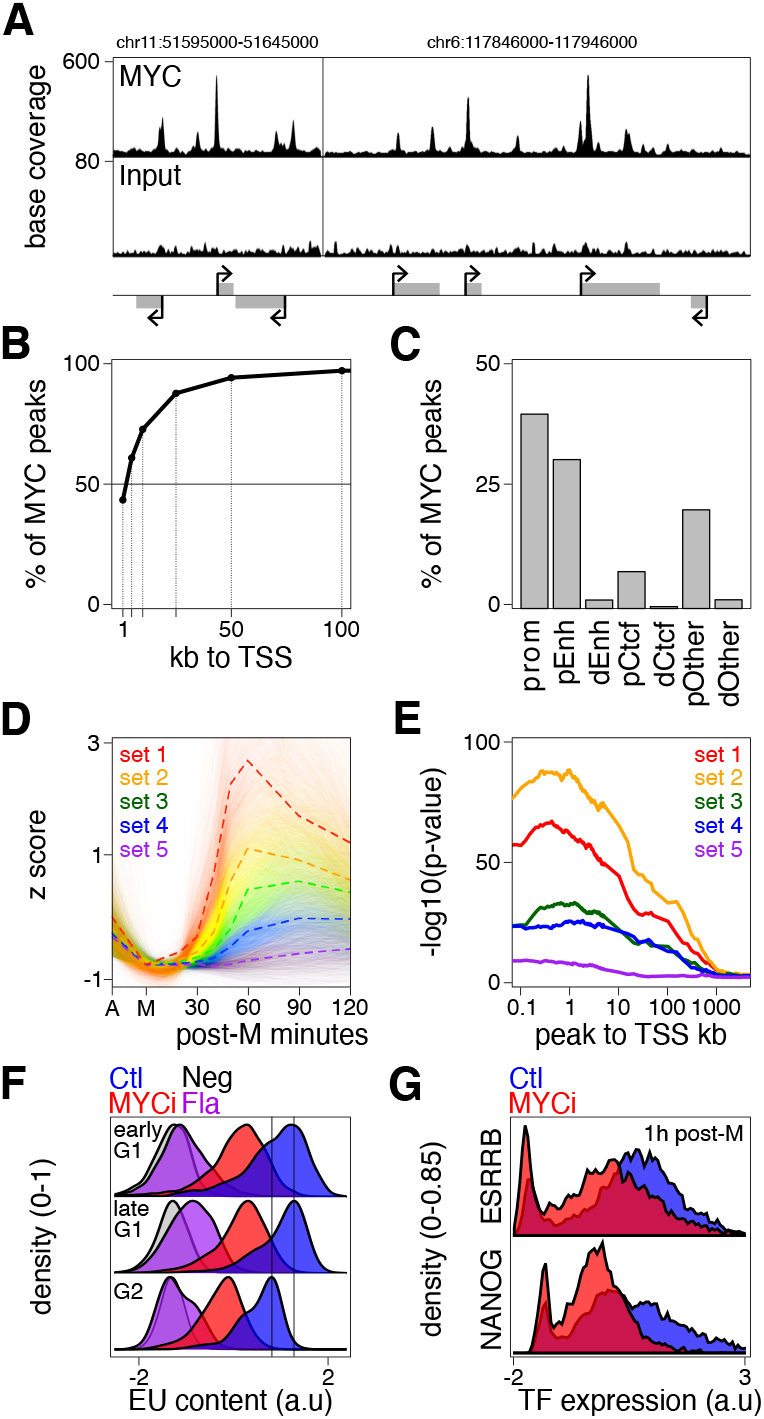
MYC drives post-mitotic ES cell hyper-transcription. **(A)** Illustrative preferential binding of MYC at promoters over two representative loci (mm10 coordinates are shown). **(B)** Cumulative proportion of MYC binding regions over increasing distances from Transcription Start Sites (TSS) showing nearly half overlap promoters and around 80-90% are located within 50kb. **(C)** Overlap of MYC binding regions with known regulatory elements such as promoters (prom), proximal (pEnh) and distal enhancers (dEnh), proximal (pCTCF) and distal CTCF binding sites (dCTCF). Binding regions not overlapping any of these categories were annotated as “Other”. Binding regions were qualified as proximal to gene TSSs when their genomic distance was below 50kb. **(D)** Relative transcriptional activity (z score) approximated by pre-mRNA quantification of around 9000 genes previously analyzed in asynchronous **(A)** and mitotic cells (M) as well as after the indicated minutes post-mitosis. A LightGBM classification algorithm was used to identify groups of genes displaying different reactivation dynamics from very fast (set 1) to very slow (set 5; see Methods for details). The plot shows individual gene traces (shadow semitransparent traces) and their corresponding mean (solid lines). **(E)** Statistical association of MYC binding sites over increasing distances from gene TSSs for the 5 sets of genes shown in (D). The plot shows the -log10(p-value) of one-sided Fisher’s exact tests. **(F)** ES cells expressing a CCNA-GFP fusion cell cycle reporter were exposed to a MYC inhibitor (MYCi) or a transcriptional inhibitor (Flavopiridol – Fla) and global transcription levels assessed by EU incorporation and FACS analysis for both early and late G1 cells, as well as G2 cells. Untreated cells (Ctl) and cells not incubated with EU (Neg) were cultured and analyzed in parallel. Two-way ANOVA revealed that there was a statistically significant interaction between the effects of the cell cycle phase and the treatment (F(6,113583) = 312.2, p <2e-16). Post hoc two-sided KS tests confirmed that the reduction of transcription observed upon MYCi treatment was statistically significant in all 3 phases (p < 2.2e-16). **(G)** ES cells were synchronized in mitosis and released into interphase in the absence (Ctl) or the presence (MYCi) of MYC inhibition. After 1h, cells were analyzed for the expression of ESRRB and NANOG by microscopy. The reduction of ESRRB and NANOG expression observed upon MYC inhibition was statistically significant (two-sided KS test p < 2.2e-16).

### MYC is not a mitotic bookmarking TF

To gain mechanistic insights into the role of MYC upon mitotic exit, we first hypothesized that it may behave as a mitotic bookmarking TF, as previously suggested in Drosophila cells^**26**^. However, we observed a nearly full depletion of MYC from mitotic chromosomes compared to a canonical mitotic bookmarking TF, ESRRB, and a striking accumulation at the mitotic spindle and centrosomes (**Fig. 2A**), as previously shown^**27**^. This was accompanied by a loss of MYC binding events from mitotic chromosomes (**Fig. 2B**,**C** and **Fig. S2A**,**B**,**C**). To gain insights in the molecular cause of the loss of MYC binding in mitosis, we analyzed protein extracts obtained from asynchronous and mitotic cells by western-blot. The expression level of MYC was reduced in mitotic cells and remained low during the release from mitosis (**Fig. 2D**). More strikingly, we observed that in interphase, MYC proteins migrate as several unresolved bands; in mitosis, however, a single high molecular weight band was observed, which reduces its apparent size as the cells reentered into interphase (**Fig. 2D**). This suggests that this TF might be, as many others^**28**^, heavily phosphorylated during mitosis. Analysis of specific phospho-MYC isophormes by immunofluorescence (**Fig. 2E**) and western-blot (**Fig. S2D**) confirmed this possibility: phosphorylation of c-MYC at Serines 62 and 373 or Threonines 58 and 358 were strongly enriched in or specific to mitotic ES cells. Since these phosphorylation events have been previously shown to control MYC stability and/or its capacity to bind DNA^**29**,**30**^, we conclude that MYC is targeted, inactivated and partially degraded by the phosphorylation cascade driving mitosis. However, rapidly after re-entry into interphase, the daughter cells dephosphorylate MYC proteins (**Fig. 2D** and **Fig. S2D**), licensing MYC for de-novo engaging in DNA binding and transcriptional activation. The question, therefore, is to understand whether, when and how is MYC recruited efficiently to its thousands of targets, particularly promoters, to trigger the reactivation of the genome. In this regard, we observed that the restoration of MYC binding starts within the first hour after mitosis (**Fig. S2E**).

**Fig. 2.**
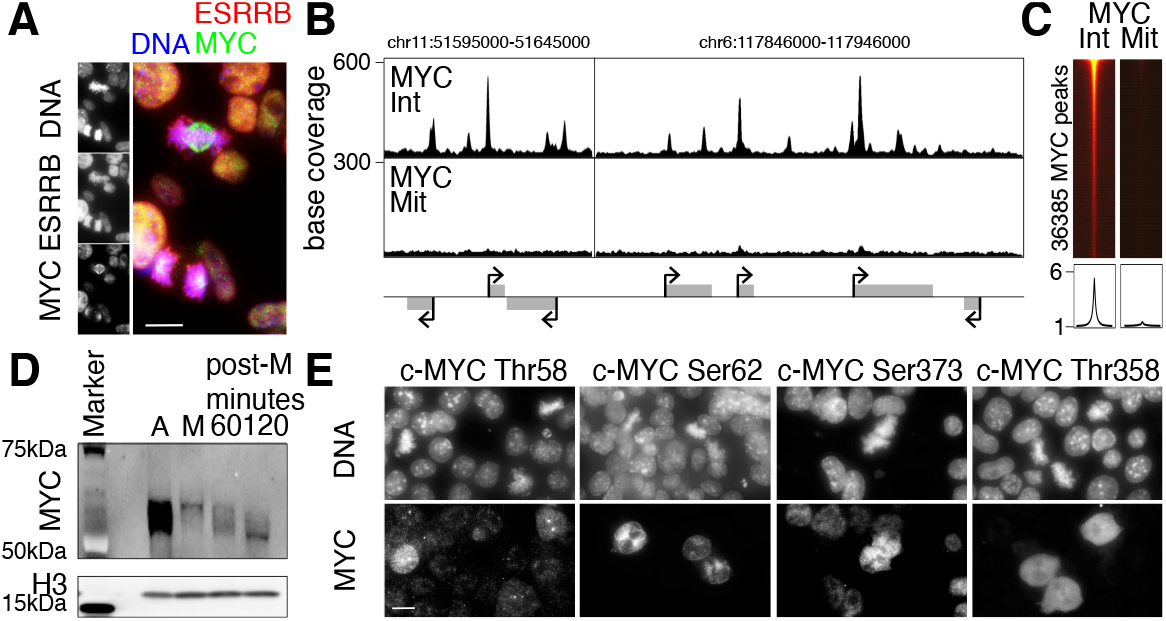
MYC is partially degraded and extensively phosphorylated in mitosis to preclude its binding to DNA. **(A)** Immuno-fluorescence of MYC and a canonical mitotic bookmarking TF (ESRRB) in ES cells. The two mitotic cells present in this representative panel (identified by DAPI staining) show an association of ESRRB but not MYC with condensed mitotic chromosomes. The horizontal white line represents 10 µm. **(B)** Illustrative loss of MYC binding during mitosis over two representative loci in interphase (Int) and during mitosis (Mit). **(C)** Direct comparison of MYC enrichment in interphase (Int) and mitosis (Mit) over 6kb-long binding regions identified in interphase and centered on the MYC binding summit (heatmaps). The plot below each heatmap displays the corresponding mean enrichment. **(D)** Western-blot analysis of MYC and a loading control (H3) in asynchronous and mitotic cells (M) as well as after the indicated minutes post-mitosis. **(E)** Immuno-fluorescence depicting four phospho-isoforms of c-MYC, as indicated above each panel. The horizontal white line represents 10 µm. Note that mitotic cells identified by DAPI staining display particularly high levels of all four phospho-isoforms.

### MAX is a mitotic bookmarking TF

In ES cells, MYC does not act as a potent mitotic bookmarking TF. Hence, we reasoned that other TFs might fulfill this role, facilitating rapid MYC binding in daughter cells. MAX, a recognized MYC interactor^**31**^, crucially guides MYC to E-boxes^**32**,**33**^, a process disrupted by the MYCi described previously^**34**^. In contrast, MAX can also homodimerize to bind DNA^**35**,**36**^. Therefore, we surmised that MAX could be behaving as a mitotic bookmarking TF of MYC targets. To test this, we first visualized MAX in live cells expressing a MAX-GFP fusion protein as they progress through mitosis (**Fig. 3A**). We observed a large and prominent retention of a MAX-GFP fusion protein on condensed mitotic chromosomes across all phases of mitosis, which we confirmed by immuno-staining of the endogenous wild-type protein (**Fig. S3A**). Moreover, we found that the binding of MAX with the mitotic chromosomes was only marginally more dynamic in mitosis than in interphase (**Fig. S3B**). Thus, during mitosis MAX behaves as other established mitotic book-marking TFs that coat the mitotic chromosomes with dynamic binding events^**28**,**37**,**38**^. We then analyzed the binding profile of MAX across mitotic chromatin (**Fig. 3B**). Among 50,612 binding sites in interphase, which are enriched for MYC binding, we found around 6,000 sites that robustly preserved MAX binding in mitosis (**Fig. 3C**,**D** and **Table S1**). These sites are enriched in promoters (**Fig. S3C**) and in the presence of E-boxes (**Fig. S3D**). Thus, at sites with high numbers of available E-boxes, MAX maintains its capacity to engage in DNA binding and can thus be considered as a canonical mitotic bookmarking TF in ES cells.

**Fig. 3.**
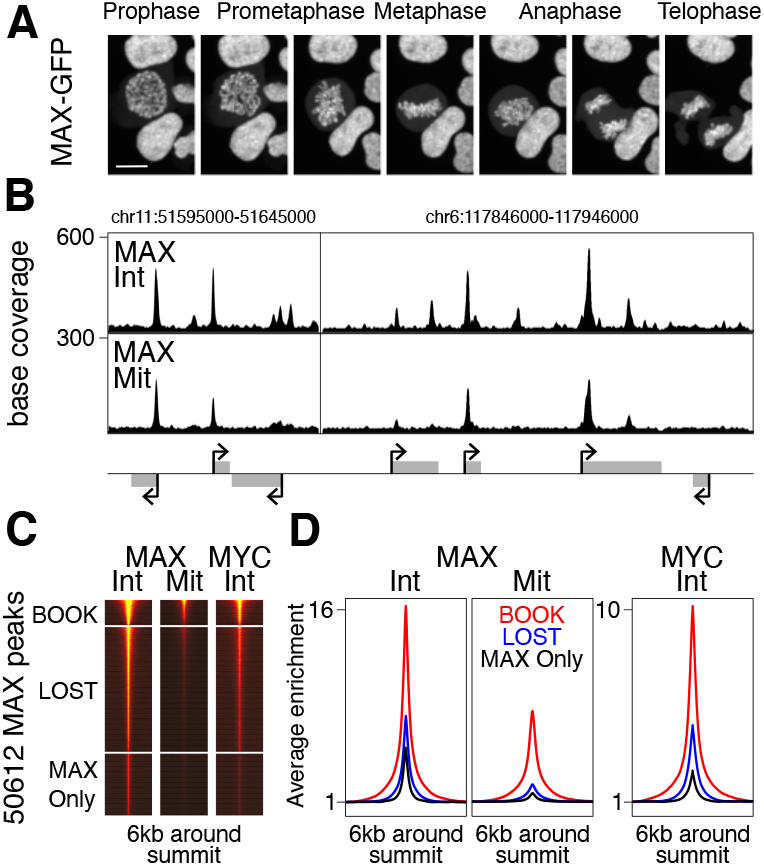
MAX is a mitotic bookmarking TF in ES cells. **(A)** Live imaging of ES cells ectopically expressing a MAX-GFP transgene, showing that MAX is strongly enriched on mitotic chromatin over all phases of mitosis, as indicated above each panel. **(B)** Illustrative mitotic bookmarking of promoters by MAX over two representative loci analyzed in interphase (Int) and mitosis (Mit). **(C)** Analysis of enrichment levels of MAX and MYC in interphase (Int) and mitosis (Mit) across all MAX binding sites clustered as subject to mitotic bookmarking (BOOK) or not (LOST, when they are associated with MYC binding and MAX only when they are not). **(D)** Average enrichment profiles of MAX and MYC binding over the categories shown in (C).

### Mitotic bookmarking by MAX is associated with efficient post-mitotic transcription

Examining the expression dynamics of genes in two illustrative loci (**Fig. 3B**), an evident trend emerged (**Fig. 4A**): genes bookmarked by MAX exhibited rapid and robust reactivation (*Hnrnpab, Nhp2, Hnrnpf, Gm38882*), whereas genes where MAX was exclusively bound during interphase were reactivated later (*Rmnd5b, Zfp239*) and genes lacking MAX binding remained not expressed (*Phykpl, Fxyd4*). To extend this correlation on a genome-wide scale, we analyzed the five sets of genes described earlier (**Fig. 1D**), revealing a powerful association between MAX bookmarking and genes displaying fast and strong reactivation dynamics (set1; **Fig. 4B** left). Conversely, sites losing MAX binding during mitosis only showed a moderate correlation with all gene groups except with those exhibiting the most rapid and strong reactivation kinetics (**Fig. 4B** middle), underscoring the specificity of the association between set1 and binding regions book-marked by MAX. Additionally, sites where MAX binding occurred exclusively during interphase and were not targeted by MYC did not exhibit any significant enrichment for any of the five gene sets analyzed (**Fig. 4B** right). The reciprocal analysis, where genes were selected based on the presence of MAX binding within 10kb of the promoter and their mean expression trend analyzed, further confirmed the relationship between MAX bookmarking, gene expression and the efficiency of post-mitotic gene reactivation (**Fig. S4A**). In conclusion, our findings demonstrate that MAX acts as a mitotic bookmarking transcription factor for a substantial fraction of MYC binding sites, particularly at promoters. These book-marked sites are closely associated with the efficiency of gene reactivation following mitosis, shedding light on the pivotal role of MAX in orchestrating post-mitotic gene expression dynamics.

**Fig. 4.**
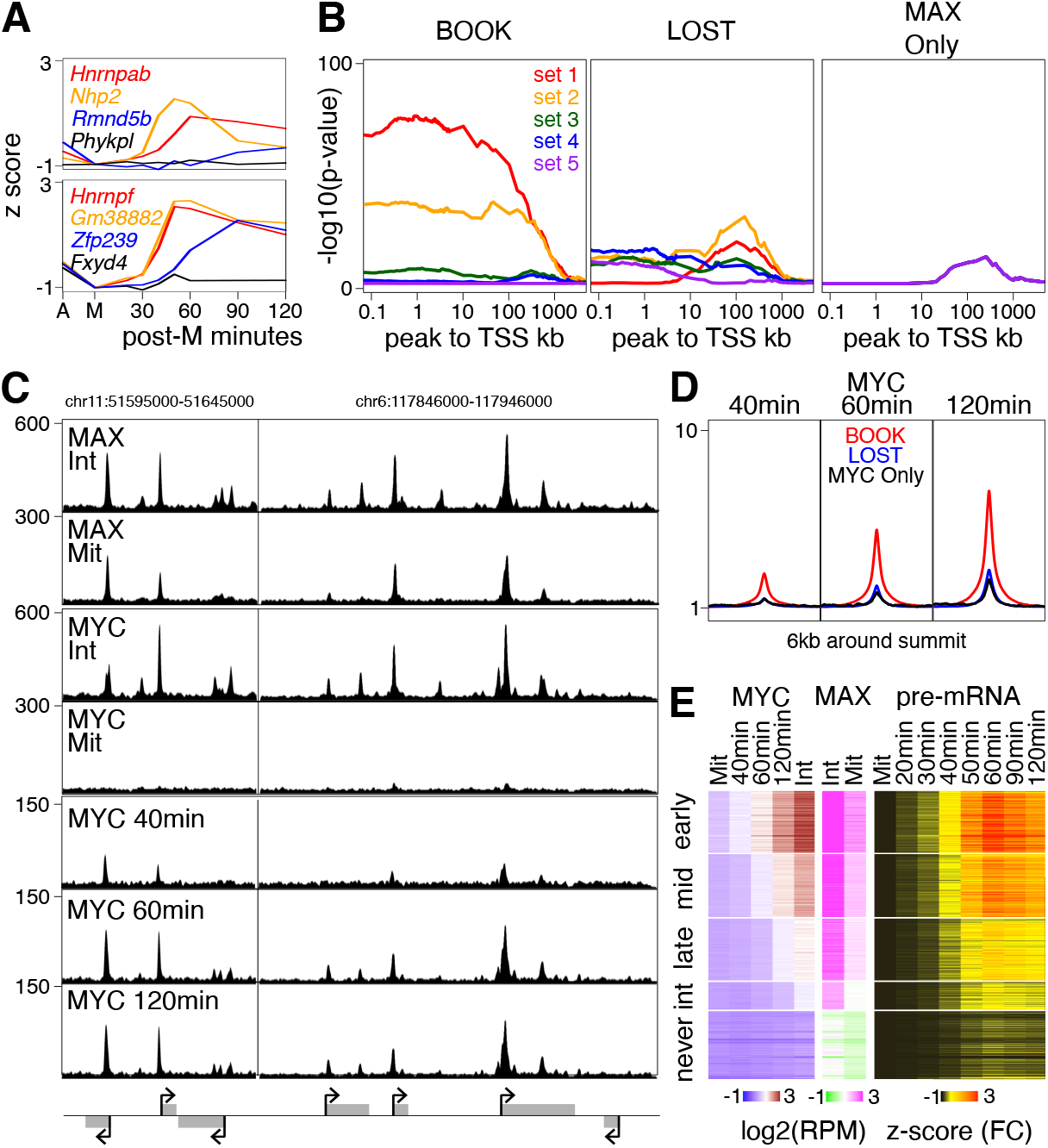
Mitotic bookmarking by MAX is associated with fast and strong post-mitotic transcription. **(A)** Transcription levels of representative gene examples subject to mitotic bookmarking (red and orange), associated with MYC/MAX binding in interphase only (blue) or not bound by MYC/MAX (black). The genes are extracted from the two representative loci shown in all figures and the plot is presented as in Figure 1D. **(B)** Statistical association of MAX binding sites over increasing distances from gene TSSs for the 5 sets of genes shown in Figure 1D, presented as in Figure 1E. Independent analyses were done for BOOK, LOST and MAX Only binding regions as described in Figure 3C. Note the exquisite specificity and strong enrichment of the most rapidly and strongly reactivated genes (sets 1 and 2) with regions subject to mitotic bookmarking by MAX. **(C)** Illustrative binding of MAX and MYC in interphase (Int) and mitosis (Mit) as well as of MYC as cells exit from mitosis (after 40, 60 and 120 minutes, as indicated), over two representative loci. **(D)** Average binding profile of MYC after 40 (left), 60 (middle) and 120 minutes post-mitosis (right) for regions subject to MAX bookmarking (BOOK) or not (LOST) as well as for MYC binding regions not associated with MAX (MYC Only). **(E)** Correlative heatmap of MYC (left) and MAX enrichment levels (middle) at all gene promoters with the reactivation of the corresponding genes (right). Gene promoters were categorized into 3 groups based on the rate of the gain of MYC binding after mitosis (early, mid, and late) plus two additional groups showing MYC enrichment exclusively in bulk interphase cells (Int) or not at all (never). Note that the groups made on the basis of MYC enrichment levels display concordant mitotic enrichments for MAX associated with fast and strong post-mitotic transcription.

### Mitotic bookmarking by MAX accelerates post-mitotic MYC binding and transcription

To further establish whether mitotic bookmarking by MAX leads to accelerated MYC recruitment, we analyzed MYC binding 40, 60 and 120 minutes after mitosis. Both for individual examples (**Fig. 4C**), and genome-wide (**Fig. 4D**), we observed that at regions subject to MAX mitotic bookmarking, MYC binding occurred earlier after mitosis compared to regions where MAX was lost. Focusing on promoters, we noticed that our conservative strategy to identify regions with robust mitotic bookmarking (see Methods) excludes promoters with potential binding events in mitosis that are associated with accelerated MYC recruitment in interphase (**Fig. S4B**). Thus, we computed MAX/MYC enrichment levels across all promoters and grouped them based on MYC enrichment levels during mitotic exit, regardless of our strict peak-calling approach (**Fig. 4E**; **Table S3**). Promoters that showed a fast gain in MYC enrichment corresponded to those strongly enriched for MAX in mitosis, resulting in more efficient postmitotic reactivation (early group in **Fig. 4E**). A gradual delay in MYC enrichment levels occurred as cells exited mitosis, correlating with a gradual reduction of mitotic MAX enrichment and with the gene reactivation strength (mid, late, int and never groups in **Fig. 4E**). To consolidate these observations, we verified that the groups of promoters inferred from direct MYC enrichment analyses displayed both coherent collective binding profiles around the transcription start site (**Fig. S4C**) and concordant associations with our selected collections of binding sites (**Fig. S4D**). Altogether, our findings highlight the importance of mitotic bookmarking by MAX in expediting MYC recruitment during the early stages of interphase, facilitating the rapid and potent reactivation of a substantial fraction of genes.

## Conclusion

Through a meticulous analysis of MYC and MAX binding dynamics, we reveal that MAX is a mitotic bookmarking TF leading to accelerated MYC recruitment during the early stages of interphase. This pivotal process facilitates the rapid and robust reactivation of a vast array of genes. Thus, our study highlights the central role of MYC/MAX in promoting efficient and global post-mitotic genome reactivation in daughter cells, a crucial feature of ES cells considering their short G1-phase^**4**,**19**^. Given the propensity of ES cells to exit self-renewal and initiate differentiation during G1^**19**,**39**^, we propose that the mechanisms described in this study place MYC/MAX as major gatekeepers of differentiation during G1. Together with other mitotic bookmarking TFs such as ESRRB and NR5A2, which display a more limited but specific effect on the reactivation of the genome^**7**^, at least two classes of mitotic bookmarking TFs preserve ES cell identity during self-renewal: on the one hand, MAX promotes strong global transcription via MYC; on the other, ESRRB, NR5A2 and other nuclear receptors provide further support and specificity to ensure the reestablishment of the pluripotency network. Since MYC/MAX are ubiquitously expressed^**40**^, it is likely that this dual scenario combining general and cell-type specific mitotic bookmarking TFs applies to other cell types. In this regard, the implications of our findings may extend into cancer biology, given the pervasive amplification and/or overexpression of MYC among cancer cells and its critical role in tumor initiation^**41**^. Understanding the regulatory mechanisms governing MYC activity is paramount in targeting the transcriptional vulnerabilities that underpin cancer development and progression^**42**,**43**^. In this context, the concept of “transcriptional addiction” – the reliance of cancer cells on sustained high transcriptional levels – has emerged as a pivotal aspect of cancer biology^**44**^. Our discovery of MAX bookmarking as a key player in MYC-driven post-mitotic hypertranscription, provides a compelling link to this concept. In the future, this may open new possibilities to disrupt the transcriptional activity that sustains the growth of cancer cells by targeting the MAX-MYC axis during the mitosis to G1-phase transition. Unraveling the complexities of gene regulation in proliferative cells, particularly across mitosis, may pave the way for novel therapeutic interventions and transformative advances in cancer treatment.

## Methods

### Cell culture and treatments

All assays were performed in E14Tg2a ES cells and derivatives, cultured on serum and LIF conditions as previously described^**16**^. Mitotic cells (>95% purity as assessed by DAPI staining and microscopy) were obtained using nocodazole shake-off^**16**^ (5h; 50 ng/ml; Sigma, M1404). For post-mitotic analyses, cells were seeded in separate dishes after the shake-off and collected after the indicated time following nocodazole withdrawal. Global transcription analyses were performed in Ccna-GFP cells^**45**^, after a short incubation (30 minutes) with EU (5-ethynyl uridine, Thermo Fisher Scientific, E10342) and in the presence or absence of a MYC inhibitor (64 µM; 10058-F4, Sigma, 475956) or of a transcription inhibitor (1 µM; Flavopiridol hydrochloride, Sigma, F3055). MAX-GFP cells were derived from E14Tg2a ES cells by stable transfection of a CAG-driven vector expressing a C-terminal fusion of MAX to GFP and linked to an IRES-Puromycin resistance cassette.

### ChIP-seq

Cells were crosslinked (50 min DSG at 2 mM; Sigma, 80424-5 mg, followed by 10 min with formaldehyde 1%; Thermo, 28908), sonicated with a Bioruptor Pico (Diagenode) and immunoprecipitated with anti-Myc (c-Myc/N-Myc (D3N8F) Rabbit), c-Myc (Thermo Fisher Scientific, PA5-85185) and anti-Max (proteintech, 10426-1-AP) antibodies. Libraries were prepared with random barcodes using NEBNext Ultra II DNA Library Prep kit for Illumina and sequenced on Illumina Illumina NextSeq 500 for 76 cycles in paired-end mode or NextSeq 2000 for 62 cycles in pairedend mode. Reads were aligned with Bowtie2^**46**^ to the mm10 genome with options “–local –very-sensitive-local –dovetail –soft-clipped-unmapped-tlen -I 0 -X 1000”.

### Protein analyses and quantifications

For Western blot, cells were lysed Laemmli buffer (BioRad, 1610737), separated using mPAGE 4-12% Bis-Tris precast gels (Merck Millipore, MP81G15) in mPAGE MOPS SDS Running Buffer (Merck Millipore, MPM0PS) and transferred onto nitrocellulose membranes (Invitrogen iBlot 2 Transfer Stacks, nitrocellulose, IB23001) using the iBlot 2 Gel Transfer Device. The membranes were incubated with primary antibodies (NANOG, Cosmobio, RCAB0001P; ESRRB, Perseus Proteomics, H6705, c-Myc/N-Myc D3N8F) and subsequently revealed with secondary antibodies (Alexa Fluor Plus 488, Goat IgG (H+L) Highly Cross-Adsorbed Secondary Antibody, anti-Rabbit and anti-mouse from Thermofisher) and a ChemiDoc MP Imaging System with Image LabTM-Touch Software Version 2.2.0.08. To analyse phosphoisoforms, proteins were extracted using a buffer containing 10 mM Tris-HCl pH 7.5, 5 mM EDTA, 150 mM NaCl, 30 mM Sodium Pyrophosphate, 50 mM Sodium Fluoride, 10% Glycerol, 1% NP40, supplemented with antiproteases (Roche Life Sciences), 2.5 U/µl Benzonase (Sigma-Aldrich), and PhosStop (Roche, 04 906 845 001). To analyse the expression of NANOG and ESRRB in single cells, 10,000 ES cells were plated on 384-well plates coated with 0.01% poly-L-ornithine (Sigma, Cat P4957) and 1X laminin (Sigma, Cat L2020), fixed with 4% formaldehyde (Thermo Scientific, 10751395) at the desired time points after mitotic release and stained first with primary antibodies (NANOG, Cosmobio, RCAB0001P; ESRRB, Perseus Proteomics, H6705) and then with secondary antibodies (Alexa-Fluor-488-conjugated, Thermo Scientific, anti-rabbit 10424752 and anti-mouse 10544773) and DAPI (BE8262). Imaging was performed using an automated spinning-disk confocal microscope Opera Phenix (PerkinElmer) with a 40X water objective. Image analyses were conducted using the software Harmony v.4.9 (Perkin Elmer).

### Global analysis of transcription

Following EU incorporation and treatment with inhibitors (see above), the cells were fixed in 4% formaldehyde in PBS and nascent RNA was labeled using the Click-iT RNA imaging kits (Thermo Fisher Scientific, C10330). Data were acquired using a FACSymphony A5 Cell Analyzer and analyzed with FlowJo software.

### Imaging

Fixation and immunofluorescence was performed with DSG+PFA and FA-only, as described, using anti-Myc (c-Myc/N-Myc (D3N8F) Rabbit), anti-Phospho-c-Myc (Thr58) (Thermo Fisher Scientific, PA5-117177), Anti-Phospho-c-Myc (Thr58) (Cell signallig, E4Z2K), anti-Phospho-c-Myc (Ser62) (Cell signallig, E1J4K), anti-Phospho-MYC-T358 (NB-22-17740-50), anti-Phospho-c-Myc (Ser373) (Invitrogen, PA537651), anti-Max (proteintech, 10426-1-AP) antibodies. Images were acquired with a Nikon Ti2E equipped with a Yokagawa CSU W1 spinning disk module and a Photometrics sCMOS Prime 95B camera using a 64× oil-immersion objective and inverted Nikon Eclipse X microscope, LUMENCOR excitation diodes, Hamamatsu ORCA-Flash 4.0LT camera and NIS Elements software. For live imaging and FRAP analyses, ES cells expressing MAX-GFP were grown on IBIDI plates, incubated with 250 nM Hoechst-33342 for 30 min before imaging and imaged at 37 °C in a humidified atmosphere (7% CO2). Images were acquired with a 63× oil immersion objective on a Nikon Ti2E equipped with a Yokagawa CSU W1 spinning disk module and a Photometrics sCMOS Prime 95B camera. For FRAP, 40 frames were acquired before bleaching (20 ms pulse using a 488nm laser, spot of minimal size) and the recovery was imaged for 1 min (1 image each 50 ms) for interphase and mitosis respectively. Fluorescence recovery was analysed in Matlab^**45**^.

### Bioinformatic and statistical analyses

Peaks were called against inputs using MACS2^**47**^. To identify different groups of peaks, we first separately identified MYC and MAX peaks (FDR < 0.01 using merged reads from either asynchronous or mitotic replicates and FDR < 0.01 in at least two individual replicates). Subsequently, we combined them all and quantified every dataset using Bamsignals package. Next, these quantifications were analyzed using a previously described generalized linear model^**16**^. Only regions displaying an enrichment versus the input with FDR < 0.05 were kept. To call a region as bookmarked it had to comply four criteria, peak calling and statistical significance of the enrichment in interphase and in mitosis. This conservative approach limits the discovery of mitotic bookmarking events but increases the robustness of the analysis. The compendium of MYC/MAX regions, their quantifications, mitotic bookmarking status and other annotations are available as Table S1. All quantifications of ChIP-seq data were systematically normalized to the library depth. To compute metaplots and enrichment heatmaps, the number of fragments covering each base of the genomic intervals under consideration were used; they were visualized with ComplexHeatmap package. Metaplots were further normalized to the minimal obtained value. Two previously described RNA-seq datasets were used for correlative purposes: one to interrogate general expression levels in undifferentiated ES cells^**48**^ and another as ES cells re-enter into interphase after a mitotic blockade^**4**^. For the latter, fold-changes after mitosis as originally reported were used after a z-score transformation for visualization purposes. This dataset was used to identify five sets of genes with distinct reactivation dynamics. For this, we built a classifier using the LightGBM algorithm^**49**^ for multi-class classification (with max_depth = 8, num_leaves = 250, learning rate = 0.01, lambda_l1: 5.05e-04, lambda_l2: 8.27e-05). Hyperparameters were tuned using a bayesian hyperparameter tuning framework optuna with 5 fold cross-validation (StratifiedKFold(n_splits=5, shuffle=True)). Number of estimators was corrected according to the early stopping option with n=100 and evaluation metric eval_metric = ‘multi_logloss’. The genes can be found in Table S2. Statistical associations between selected groups of genes and genomic intervals were assessed with one-sided Fisher’s exact tests where increasing distances from the transcription start sites were used to probe the effect of the distance from 100bp to 5Mb, as described^**50**^. The genome-wide location of Myc/Max motifs (E-boxes) was downloaded from the Jaspar database and only motifs with p<0.001 obtained from the 4 available predictions for slightly different motifs (MA0058.3, MA0059.1, MA0104.4, MA0147.3) were combined and used to compute the number of motifs per ChIP-seq peak. To control for peak width variability, we systematically considered 1kb windows centered on the generic peak summit obtained from the average of MYC and MAX coverage. To classify binding regions as promoters, enhancers or CTCF binding sites, we used the 5’ end region of all known genes and collections of known regulatory elements^**15**,**51**^. All statistical tests used are reported in the legends of the figures.

## Acknowledgements

The authors acknowledge the Biomics, Flow Cytometry, Photonic BioImaging and Image Analysis Hub platforms of Institut Pasteur. PN acknowledges the Institut Pasteur, the CNRS, and Revive (Investissement d’Avenir; ANR-10-LABX-73) for recurrent funding and the Agence Nationale de la Recherche (ANR20CE12002801 CHRODYNE), Ligue contre le Cancer (LNCC EL2018 NAVARRO) and the European Research Council (ERC-CoG-2017 BIND) for financial support.

## Author contributions

IG performed all experiments and preliminary bioinformatic analyses with help from LAP. AC performed final bioinformatic and statistical analyses. PE, FM and AD provided essential help with FACS/OPERA, FRAP and cell culture, respectively. PN conceived the project, analyzed the data and wrote the manuscript with IG.

## Declaration of interests

The authors declare no competing interests.

## Supplementary information

Four supplementary figures accompany this manuscript, they can be found at the end of this document. Three Supplementary Tables are available online.

---

**Supplementary Information, Fig. S 1.**
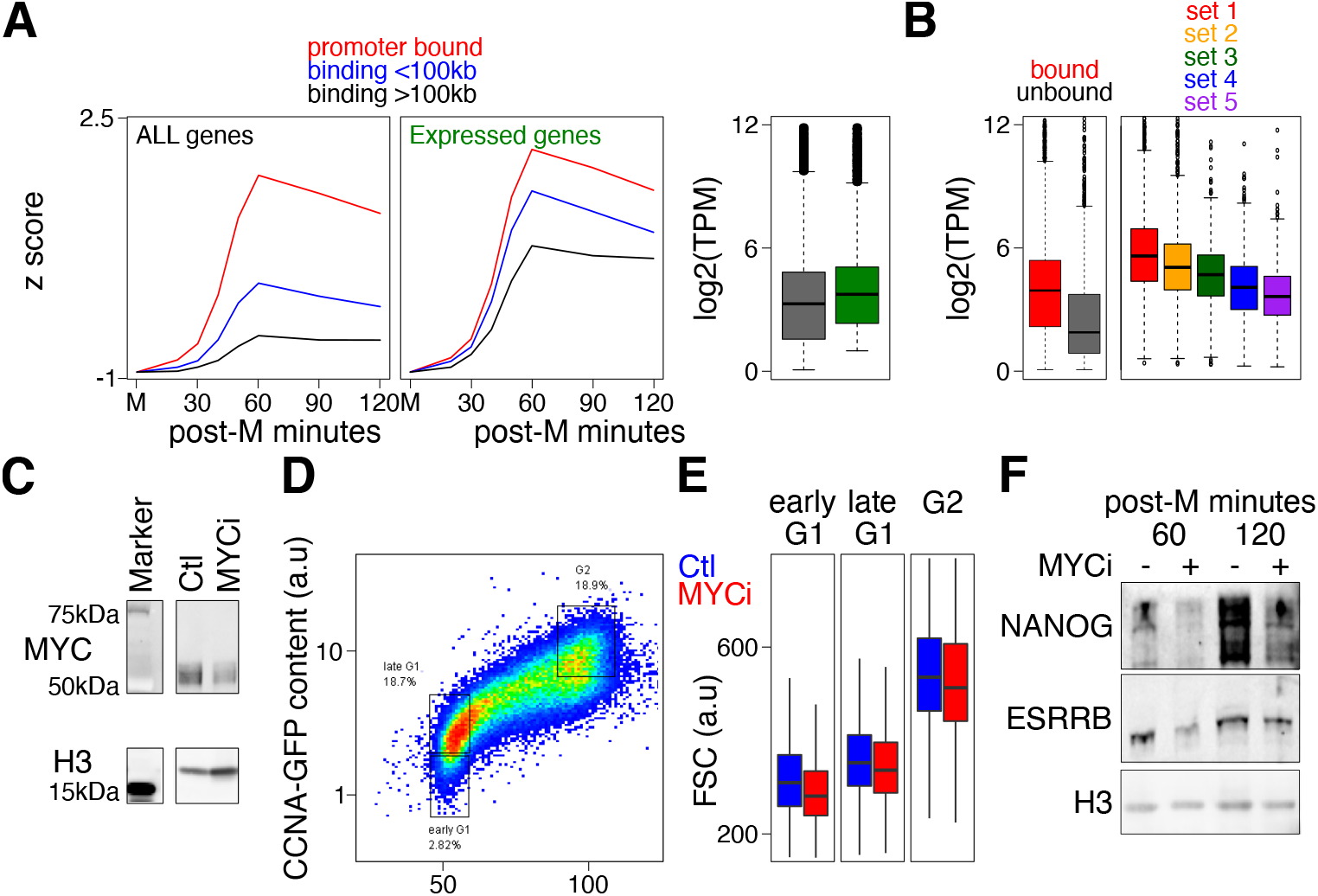
Additional information on MYC association with post-mitotic hyper-transcription. **(A)** Mean gene reactivation presented as in Figure 1D for either all (left) or expressed genes only (middle) for groups of genes displaying MYC binding at the promoter (red), within 100kb (blue) or at more than 100kb (black) from the TSS. The boxplot (median – bar; 25-75% percentiles – box; 1.5-folds the inter-quartile range – whiskers) on the right shows the expression level (Transcripts per million; TPM) of all genes (gray) or of the selection of expressed genes (green). **(B)** Boxplots of the expression levels of genes bound or not by MYC at the promoter (left) or belonging to the 5 sets of genes displaying distinct reactivation kinetics after mitosis, presented as in (A). **(C)** Western-blot of MYC and a loading control (H3) in untreated (Ctl) or in cells treated with a MYC inhibitor (MYCi). **(D)** FACS analysis of CCNA-GFP cells displaying CCNA-GFP levels (Y-axis) and DNA content (X-axis) as well as the gates used to subset early G1, late G1 and G2 cells in the analysis of global transcription assessed by EU incorporation in Figure 1F. **(E)** Analysis of the size of the cells by FACS (FSC parameter) upon MYC inhibition (MYCi). Two-way ANOVA revealed that there was a statistically significant interaction between the effects of the cell cycle phase and the treatment (F(6,113583) = 59.44, p <2e-16). Post hoc two-sided KS tests confirmed that the reduction of cell size was statistically significant upon MYCi treatment in all 3 phases (p < 2.2e-16). **(F)** Western-blot analysis of NANOG and ESRRB confirms the decrease in their expression level observed 60 and 120 min after mitosis in the presence of MYC inhibitors

**Supplementary Information, Fig. S 2.**
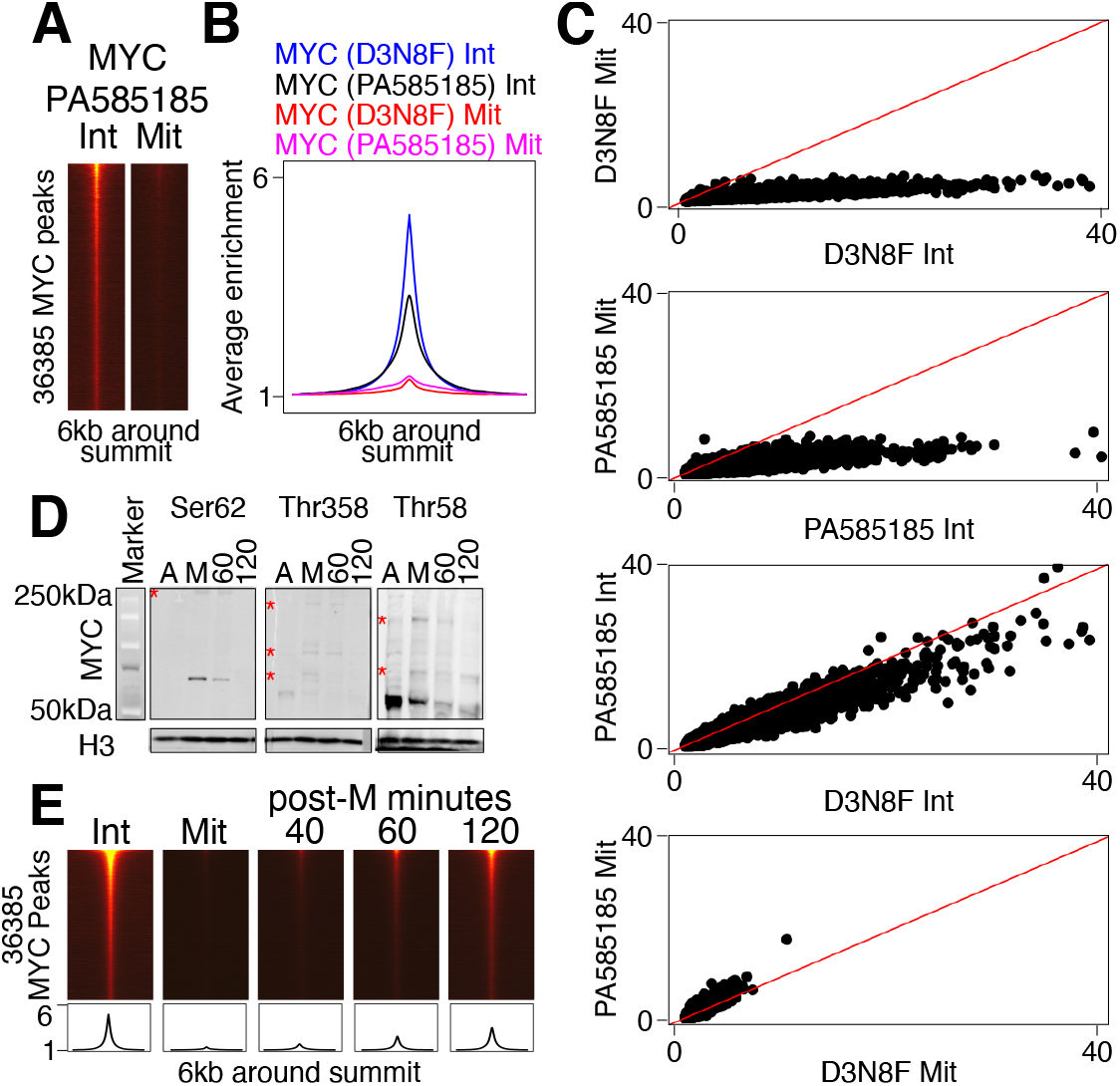
Additional information on MYC behavior during mitosis. **(A)** Enrichment levels of MYC in interphase and mitosis using an independent antibody specific of c-MYC, presented as in Figure 2C. **(B)** Average enrichment profiles of MYC binding using generic MYC antibodies (D3N8F) or c-MYC specific (PA585185) in interphase and in mitosis. **(C)** Scatter-plots showing pair-wise comparisons between antibodies and samples (interphase or mitosis). Y and X-axis show fragments per million of the indicated quantification. The red line shows the 1:1 ratio. **(D)** Western-blot analysis of three c-MYC phospho-isoforms in asynchronous (A) or mitotic cells (M), as well as 60 or 120 minutes post-mitosis. The red asterisks denote bands of particularly high molecular weight likely reflecting additional post-translational modifications, possibly ubiquitination to target phospho-MYC for degradation. **(E)** Heatmaps displaying MYC enrichment levels as cells exit from mitosis (40, 60 and 120 minutes post-mitosis), with the corresponding average plot show below. Note that MYC binding is licensed after 40-60 minutes, albeit at low levels.

**Supplementary Information, Fig. S 3.**
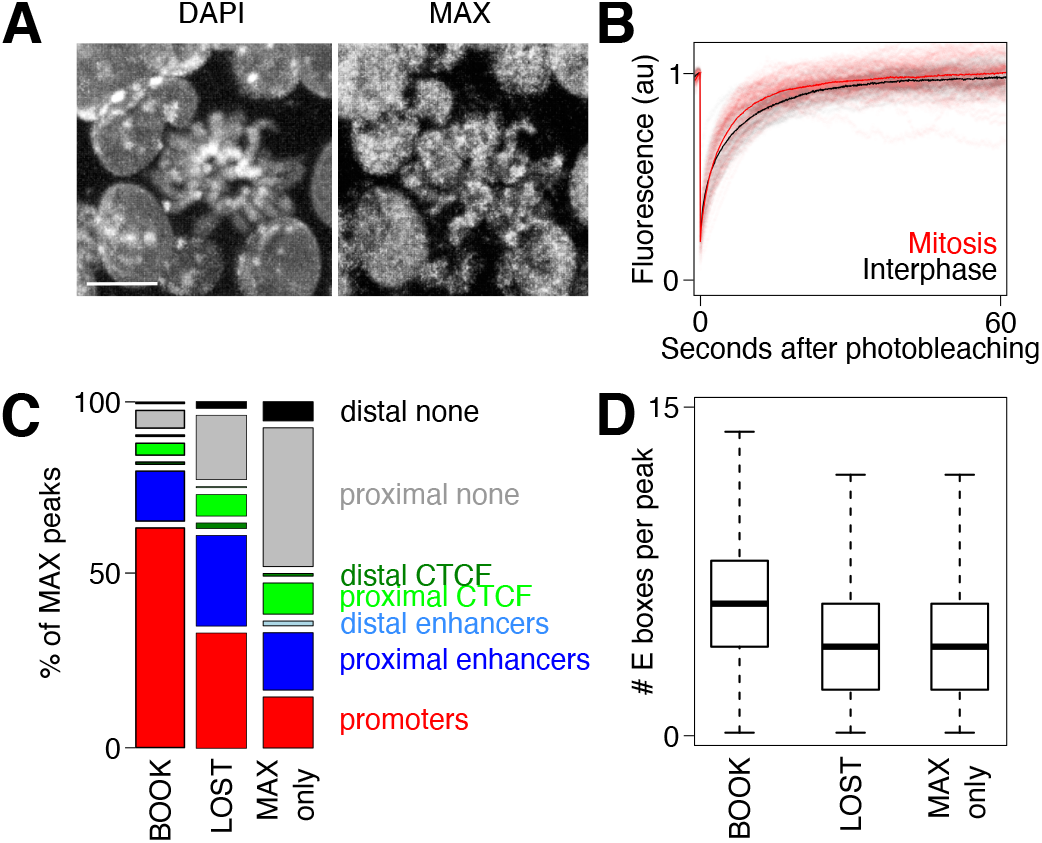
Additional information on MAX behavior during mitosis. **(A)** Immuno-fluorescence of endogenously expressed MAX displaying a strong enrichment on mitotic chromosomes. The horizontal white line represents 10 µm. **(B)** Results of FRAP assays showing a slightly accelerated recovery of MAX-GFP signal on mitotic chromosomes (red) compared to interphase (black). The plot shows individual measurements (46 cells in interphase and 92 in mitosis from 3 independent experiments; shadow semitransparent traces) and their corresponding mean (solid lines). **(C)** Distribution of MAX peaks categorized as subject to mitotic bookmarking (BOOK) or not (LOST and MAX Only), as shown in Figure 3C, and associated with different gene regulatory elements listed on the right (established as in Figure 1C). Pearson’s Chi-squared test revealed a statistically significant association between binding status and the type of region (X2(12, 50612) = 8768.8, p < 2.2e-16). Subsequent Fisher’s exact test showed the enrichment of promoters at MAX bookmarked regions was statistically significant (p < 2.2e-16; odds ratio = 5.85). **(D)** Boxplot (median – bar; 25-75% percentiles – box; 1.5-folds the inter-quartile range – whiskers) showing the number of E-boxes identified at 1kb-long regions centered on the summit of MAX binding regions for the three categories of regions identified for MAX in Figure 3C. Pearson’s Chi-squared test indicated a statistically significant association between the number of motifs and the type of region (X2(108, 51348) = 2763.8, p < 2.2e-16). Mann-Whitney tests confirmed the statistical significance of the enrichment of motifs at bookmarked regions versus lost and MAX only regions (p < 2.2e-16).

**Supplementary Information, Fig. S 4.**
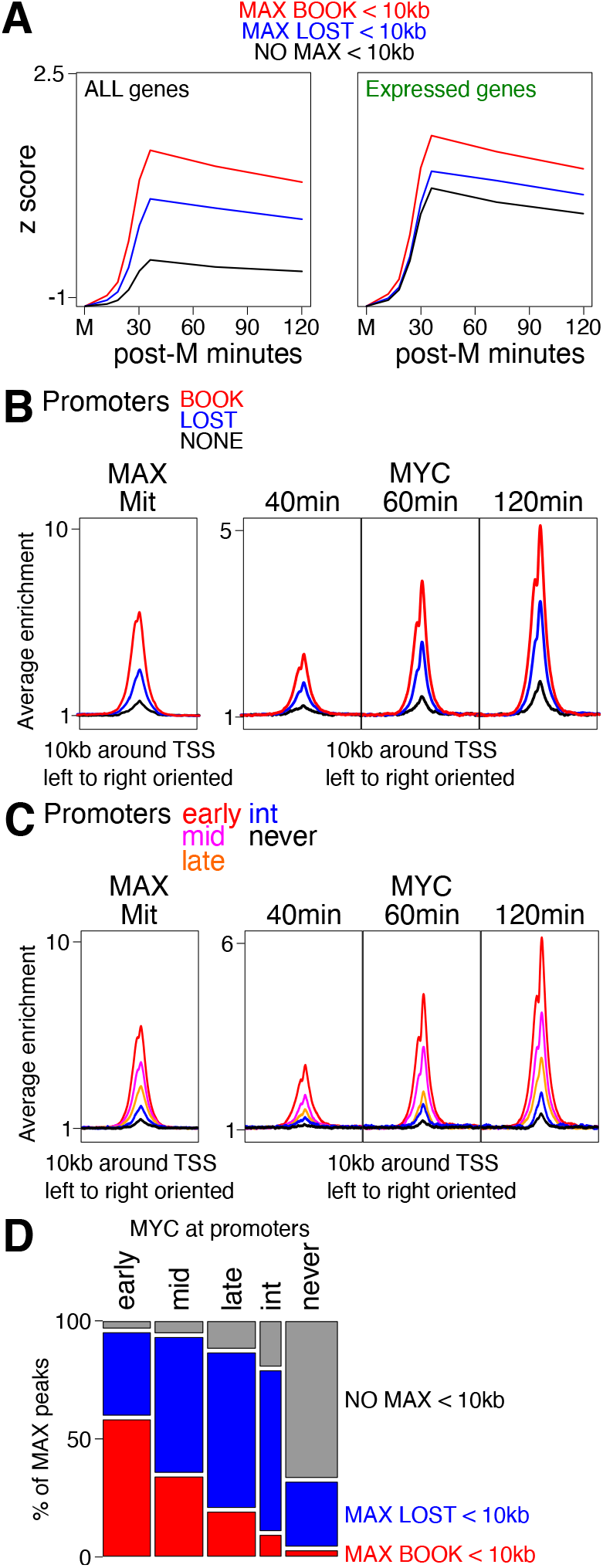
Additional information on MYC/MAX association with post-mitotic gene reactivation. **(A)** Mean gene reactivation presented as in Figure 1D for either all (left) or expressed genes only (right) for groups of genes displaying different categories of MAX binding regions within 10kb of the TSS, as indicated. **(B)** Average enrichment profile of MAX in mitosis (Mit) and of MYC after 40 (left), 60 (middle) and 120 minutes post-mitosis (right). Ten kilobases-long regions centered on the TSS of all genes are shown. **(C)** Identical analysis to (B) but for promoters displaying different dynamics of MYC binding after mitosis, as shown in Figure 4E. **(D)** Distribution of MAX peak categories (listed on the right) across the promoter groups identified in Figure 4E (listed on top). Pearson’s Chi-squared test indicated a statistically significant association between promoter categories and MAX binding categories (X2(8, 16012) = 9486.28, p < 2.2e-16). Subsequent Fisher’s exact test showed that the enrichment of bookmarked regions at early promoters was statistically significant (p < 2.2e-16; odds ratio = 6.83).

## Notes

### Competing Interest Statement

The authors have declared no competing interest.

## References

1. Hsiung CC, Bartman CR, Huang P, Ginart P, Stonestrom AJ, Keller CA, Face C, Jahn KS, Evans P, Sankaranarayanan L, Giardine B, Hardison RC, Raj A, Blobel GA. A hyperactive transcriptional state marks genome reactivation at the mitosis-G1 transition. Genes Dev. 2016 Jun 15;30(12):1423–39.

2. Palozola KC, Liu H, Nicetto D, Zaret KS. Low-Level, Global Transcription during Mitosis and Dynamic Gene Reactivation during Mitotic Exit. Cold Spring Harb Symp Quant Biol. 2017;82:197–205.

3. Kang H, Shokhirev MN, Xu Z, Chandran S, Dixon JR, Hetzer MW. Dynamic regulation of histone modifications and long-range chromosomal interactions during postmitotic transcriptional reactivation. Genes Dev. 2020 Jul 1;34(13-14):913–930.

4. Chervova A, Festuccia N, Altamirano-Pacheco L, Dubois A, Navarro P. A gene subset requires CTCF bookmarking during the fast post-mitotic reactivation of mouse ES cells. EMBO Rep. 2023 Jan 9;24(1):e56075.

5. Pelham-Webb B, Polyzos A, Wojenski L, Kloetgen A, Li J, Di Giammartino DC, Sakellaropoulos T, Tsirigos A, Core L, Apostolou E. H3K27ac bookmarking promotes rapid post-mitotic activation of the pluripotent stem cell program without impacting 3D chromatin reorganization. Mol Cell. 2021 Apr 15;81(8):1732–1748.e8.

6. Zhu Z, Chen X, Guo A, Manzano T, Walsh PJ, Wills KM, Halliburton R, Radko-Juettner S, Carter RD, Partridge JF, Green DR, Zhang J, Roberts CWM. Mitotic bookmarking by SWI/SNF subunits. Nature. 2023 Jun;618(7963):180–187.

7. Almira Chervova, Amandine Molliex, H. Irem Baymaz, Thaleia Papadopoulou, Florian Mueller, Eslande Hercul, David Fournier, Agnès Dubois, Nicolas Gaiani, Petra Beli, Nicola Festuccia, Pablo Navarro. Mitotic bookmarking redundancy by nuclear receptors mediates robust postmitotic reactivation of the pluripotency network. 2022 Biorxiv preprint https://doi.org/10.1101/2022.11.28.518105

8. Silja Placzek, Ludovica Vanzan, David M Suter. Orchestration of pluripotent stem cell genome reactivation during mitotic exit. 2023 Biorxiv preprint https://doi.org/10.1101/2023.08.09.552605

9. Wiman KG, Zhivotovsky B. Understanding cell cycle and cell death regulation provides novel weapons against human diseases. J Intern Med. 2017 May;281(5):483–495

10. Madden SK, de Araujo AD, Gerhardt M, Fairlie DP, Mason JM. Taking the Myc out of cancer: toward therapeutic strategies to directly inhibit c-Myc. Mol Cancer. 2021 Jan 4;20(1):3.

11. Wang C, Zhang J, Yin J, Gan Y, Xu S, Gu Y, Huang W. Alternative approaches to target Myc for cancer treatment. Signal Transduct Target Ther. 2021 Mar 10;6(1):117.

12. Llombart V, Mansour MR. Therapeutic targeting of “undruggable” MYC. EBioMedicine. 2022 Jan;75:103756.

13. Gottesfeld JM, Forbes DJ. Mitotic repression of the transcriptional machinery. Trends Biochem Sci. 1997 Jun;22(6):197–202.

14. Gonzalez I, Molliex A, Navarro P. Mitotic memories of gene activity. Curr Opin Cell Biol. 2021 Apr;69:41–47.

15. Ito K, Zaret KS. Maintaining Transcriptional Specificity Through Mitosis. Annu Rev Genomics Hum Genet. 2022 Aug 31;23:53–71.

16. Festuccia N, Owens N, Papadopoulou T, Gonzalez I, Tachtsidi A, Vandoermel-Pournin S, Gallego E, Gutierrez N, Dubois A, Cohen-Tannoudji M, Navarro P. Transcription factor activity and nucleosome organization in mitosis. Genome Res. 2019 Feb;29(2):250–260.

17. Price RM, Budzyński MA, Shen J, Mitchell JE, Kwan JZJ, Teves SS. Heat shock transcription factors demonstrate a distinct mode of interaction with mitotic chromosomes. Nucleic Acids Res. 2023 Jun 9;51(10):5040–5055.

18. Teves SS, An L, Bhargava-Shah A, Xie L, Darzacq X, Tjian R. A stable mode of bookmarking by TBP recruits RNA polymerase II to mitotic chromosomes. Elife. 2018 Jun 25;7:e35621.

19. Coronado D, Godet M, Bourillot PY, Tapponnier Y, Bernat A, Petit M, Afanassieff M, Markossian S, Malashicheva A, Iacone R, Anastassiadis K, Savatier P. A short G1 phase is an intrinsic determinant of naïve embryonic stem cell pluripotency. Stem Cell Res. 2013 Jan;10(1):118–31.

20. Nie Z, Hu G, Wei G, Cui K, Yamane A, Resch W, Wang R, Green DR, Tessarollo L, Casellas R, Zhao K, Levens D. c-Myc is a universal amplifier of expressed genes in lymphocytes and embryonic stem cells. Cell. 2012 Sep 28;151(1):68–79.

21. Lin CY, Lovén J, Rahl PB, Paranal RM, Burge CB, Bradner JE, Lee TI, Young RA. Transcriptional amplification in tumor cells with elevated c-Myc. Cell. 2012 Sep 28;151(1):56–67.

22. Percharde M, Wong P, Ramalho-Santos M. Global Hypertranscription in the Mouse Embryonic Germline. Cell Rep. 2017 Jun 6;19(10):1987–1996.

23. Lorenzin F, Benary U, Baluapuri A, Walz S, Jung LA, von Eyss B, Kisker C, Wolf J, Eilers M, Wolf E. Different promoter affinities account for specificity in MYC-dependent gene regulation. Elife. 2016 Jul 27;5:e15161.

24. Kim J, Chu J, Shen X, Wang J, Orkin SH. An extended transcriptional network for pluripotency of embryonic stem cells. Cell. 2008 Mar 21;132(6):1049–61.

25. Kim J, Woo AJ, Chu J, Snow JW, Fujiwara Y, Kim CG, Cantor AB, Orkin SH. A Myc network accounts for similarities between embryonic stem and cancer cell transcription programs. Cell. 2010 Oct 15;143(2):313–24.

26. Yang J, Sung E, Donlin-Asp PG, Corces VG. A subset of Drosophila Myc sites remain associated with mitotic chromosomes colocalized with insulator proteins. Nat Commun. 2013;4:1464.

27. Becker S, Kiecke C, Schäfer E, Sinzig U, Deuper L, Trigo-Mourino P, Griesinger C, Koch R, Rydzynska Z, Chapuy B, von Bonin F, Kube D, Venkataramani V, Bohnenberger H, Leha A, Flach J, Dierks S, Bastians H, Maruschak B, Bojarczuk K, Taveira MO, Trümper L, Wulf GM, Wulf GG. Destruction of a Microtubule-Bound MYC Reservoir during Mitosis Contributes to Vincristine’s Anticancer Activity. Mol Cancer Res. 2020 Jun;18(6):859–872.

28. Festuccia N, Gonzalez I, Owens N, Navarro P. Mitotic bookmarking in development and stem cells. Development. 2017 Oct 15;144(20):3633–3645.

29. Huang Z, Traugh JA, Bishop JM. Negative control of the Myc protein by the stress-responsive kinase Pak2. Mol Cell Biol. 2004 Feb;24(4):1582–94.

30. Macek P, Cliff MJ, Embrey KJ, Holdgate GA, Nissink JWM, Panova S, Waltho JP, Davies RA. Myc phosphorylation in its basic helix-loop-helix region destabilizes transient α-helical structures, disrupting Max and DNA binding. J Biol Chem. 2018 Jun 15;293(24):9301–9310.

31. Blackwood EM, Lüscher B, Eisenman RN. Myc and Max associate in vivo. Genes Dev. 1992 Jan;6(1):71–80.

32. Blackwell TK, Kretzner L, Blackwood EM, Eisenman RN, Weintraub H. Sequence-specific DNA binding by the c-Myc protein. Science. 1990 Nov 23;250(4984):1149–51.

33. Walhout AJ, Gubbels JM, Bernards R, van der Vliet PC, Timmers HT. c-Myc/Max heterodimers bind cooperatively to the E-box sequences located in the first intron of the rat ornithine decarboxylase (ODC) gene. Nucleic Acids Res. 1997 Apr 15;25(8):1493–501.

34. Yin X, Giap C, Lazo JS, Prochownik EV. Low molecular weight inhibitors of Myc-Max interaction and function. Oncogene. 2003 Sep 18;22(40):6151–9.

35. Ferré-D’Amaré AR, Prendergast GC, Ziff EB, Burley SK. Recognition by Max of its cognate DNA through a dimeric b/HLH/Z domain. Nature. 1993 May 6;363(6424):38–45.

36. Brownlie P, Ceska T, Lamers M, Romier C, Stier G, Teo H, Suck D. The crystal structure of an intact human Max-DNA complex: new insights into mechanisms of transcriptional control. Structure. 1997 Apr 15;5(4):509–20.

37. Teves SS, An L, Hansen AS, Xie L, Darzacq X, Tjian R. A dynamic mode of mitotic bookmarking by transcription factors. Elife. 2016 Nov 19;5:e22280.

38. Raccaud M, Friman ET, Alber AB, Agarwal H, Deluz C, Kuhn T, Gebhardt JCM, Suter DM. Mitotic chromosome binding predicts transcription factor properties in interphase. Nat Commun. 2019 Jan 30;10(1):487.

39. Pauklin S, Vallier L. The Cell-Cycle State of Stem Cells Determines Cell Fate Propensity. Cell. 2014 Mar 13;156(6):1338.

40. Cascón A, Robledo M. MAX and MYC: a heritable breakup. Cancer Res. 2012 Jul 1;72(13):3119–24.

41. Gabay M, Li Y, Felsher DW. MYC activation is a hallmark of cancer initiation and maintenance. Cold Spring Harb Perspect Med. 2014 Jun 2;4(6):a014241.

42. Bushweller JH. Targeting transcription factors in cancer - from undruggable to reality. Nat Rev Cancer. 2019 Nov;19(11):611–624.

43. Vervoort SJ, Devlin JR, Kwiatkowski N, Teng M, Gray NS, Johnstone RW. Targeting transcription cycles in cancer. Nat Rev Cancer. 2022 Jan;22(1):5–24.

44. Bradner JE, Hnisz D, Young RA. Transcriptional Addiction in Cancer. Cell. 2017 Feb 9;168(4):629–643.

45. Festuccia N, Dubois A, Vandormael-Pournin S, Gallego Tejeda E, Mouren A, Bessonnard S, Mueller F, Proux C, Cohen-Tannoudji M, Navarro P. Mitotic binding of Esrrb marks key regulatory regions of the pluripotency network. Nat Cell Biol. 2016 Nov;18(11):1139–1148.

46. Langmead B, Salzberg SL. Fast gapped-read alignment with Bowtie 2. Nat Methods. 2012 Mar 4;9(4):357–9.

47. Feng J, Liu T, Qin B, Zhang Y, Liu XS. Identifying ChIP-seq enrichment using MACS. Nat Protoc. 2012 Sep;7(9):1728–40.

48. Dubois A, Vincenti L, Chervova A, Greenberg MVC, Vandormael-Pournin S, Bourc’his D, Cohen-Tannoudji M, Navarro P. H3K9 trimethylation at Nanog times differentiation commitment and enables the acquisition of primitive endoderm fate. Development. 2022 Sep 1;149(17):dev201074.

49. Ke, G., Meng, Q., Finley, T., Wang, T., Chen, W., Ma, W., Ye, Q. and Liu, T.Y., 2017. Lightgbm: A highly efficient gradient boosting decision tree. Advances in neural information processing systems, 30.

50. Heurtier V, Owens N, Gonzalez I, Mueller F, Proux C, Mornico D, Clerc P, Dubois A, Navarro P. The molecular logic of Nanog-induced self-renewal in mouse embryonic stem cells. Nat Commun. 2019 Mar 7;10(1):1109.

51. González-Ramírez M, Ballaré C, Mugianesi F, Beringer M, Santanach A, Blanco E, Di Croce L. Differential contribution to gene expression prediction of histone modifications at enhancers or promoters. PLoS Comput Biol. 2021 Sep 2;17(9):e1009368.

